# SEGUID v2: Extending SEGUID checksums for circular, linear, single- and double-stranded biological sequences

**DOI:** 10.1101/2024.02.28.582384

**Authors:** Humberto Pereira, Paulo César Silva, M. Wayne Davis, Louis Abraham, György Babnigg, Henrik Bengtsson, Björn Johansson

**Affiliations:** Center of Molecular and Environmental Biology Engineering, University of Minho, Campus de Gualtar, Braga, 4710-057, Portugal; Howard Hughes Medical Institute and School of Biological Sciences, University of Utah, Salt Lake City, UT, USA; PRISM, Université Paris 1, Panthéon–Sorbonne, Paris, France; Biosciences Division, Argonne National Laboratory, Argonne, IL, USA; Department of Epidemiology and Biostatistics, University of California, San Francisco, CA, USA; Helen Diller Family Comprehensive Cancer Center, University of California, San Francisco, CA, USA

**Keywords:** checksum, hash, DNA, RNA, protein, SHA-1, Base64url, SEGUID

## Abstract

**Background:** Synthetic biology involves combining different DNA fragments, each containing functional biological parts, to address specific problems. Fundamental gene-function research often requires cloning and propagating DNA fragments, such as those from the iGEM Parts Registry or Addgene, typically distributed as circular plasmids. Addgene’s repository alone offers around 150,000 plasmids. To ensure data integrity, cryptographic checksums can be calculated for the sequences. Each sequence has a unique checksum, making checksums useful for validation and quick lookups of associated annotations. For example, the SEGUID checksum uniquely identifies protein sequences with a 27-character string.

**Objectives:** The original SEGUID, while effective for protein sequences and single-stranded DNA (ssDNA), is not suitable for circular DNA since there is no natural starting position nor for double-stranded DNA (dsDNA) since two separate sequences are present. Challenges include how to uniquely represent linear dsDNA, circular ssDNA, and circular dsDNA. To meet these needs, we propose SEGUID v2, which extends the original SEGUID to handle additional types of sequences.

**Conclusions:** SEGUID v2 produces orientation and rotation invariant checksums for single-stranded, double-stranded, possibly staggered, linear, and circular DNA and RNA sequences. Customizable alphabets allow for other types of sequences. In contrast to the original SEGUID, which uses Base64, SEGUID v2 uses Base64url to encode the SHA-1 hash. This ensures SEGUID v2 checksums can be used as-is in filenames, regardless of platform, and in URLs, with minimal friction.

**Availability:** SEGUID v2 is readily available for major programming languages, distributed under the MIT license. JavaScript package **seguid** is available on npm, Python package **seguid** on PyPi, R package **seguid** on CRAN, and a Tcl script on GitHub. These tools, along with documentation, examples, and an online **SEGUID Calculator**, can be found at https://www.seguid.org.

## 1 Introduction

The post-genomic era has provided access to more biological sequence data than ever before. These sequence data provide an important resource for both fundamental research and synthetic biology, which aim to understand and harness biology to solve urgent societal problems, such as global warming or food security. Genetic parts contained on DNA fragments are distributed to and from culture collections and repositories, as well as within the research community to study genes and build new synthetic systems. These parts are almost always distributed as plasmids, where the DNA of interest is inserted into a circular double-stranded (ds) DNA plasmid providing replication and selection in a microorganism optimal for producing high-quality DNA, usually *E. coli*.

Each plasmid should have an associated sequence file that is distributed along with the physical clone. Although repositories such as Addgene (Kamens, 2014b,a) and the iGEM (International Genetically Engineered Machine) Parts collection provide standards, the transmission of biological material and associated information is still very much an ad-hoc procedure.

Data integrity is of great importance since it directly affects the reproducibility of science that relies on the original genetic parts. Checksums are widely used for verifying that information is preserved through space and time, and have also been applied to safeguard biological sequences. A checksum is a short string of characters that is calculated from, and typically stored with, arbitrarily sized data. By transmitting the checksum together with the data, a receiver can verify the correctness of the data. Checksums are widely used, but not directly applicable due to DNA topology. SEGUID v2, which is a topology-aware extension to SEGUID v1, addresses this limitation.

Several existing checksum algorithms are relevant in bioinformatics. One of the first used was the cyclic redundancy check (**CRC**) (W.H. et al., 1993), which is a computationally inexpensive, error-detecting algorithm originally designed for network and storage devices. The UniProt database initially used a 64-bit **CRC** (**CRC-64**) checksum^1^. This is the reason why the UniProt Archive (UniParc) still provides **CRC-64**^2^. A disadvantage of the **CRC-64** is that collisions have been observed for biological data. For example, the two protein sequences with GenBank accession numbers *AAF20475*.*1*_GB_^3^ and *AAF20481*.*1*_GB_^4^, have the same **CRC-64** hexadecimal checksum, despite differing in two of the 110 amino acids (Figure 1).

**Figure 1:**
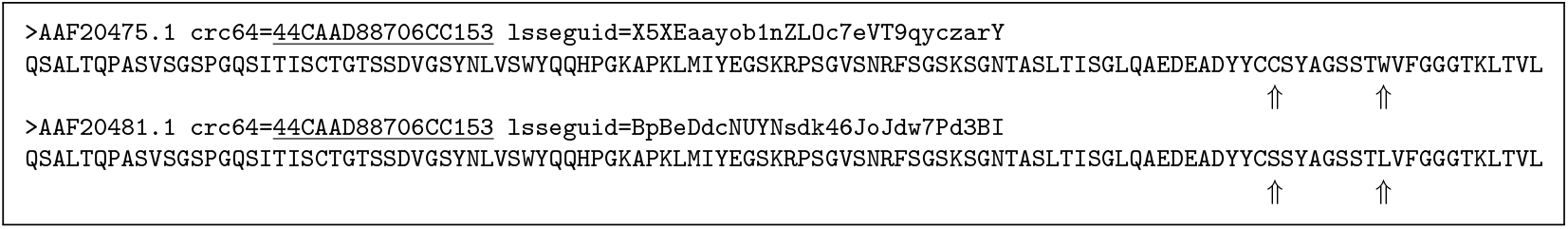
Two protein sequences with colliding CRC-64 checksums. The two protein sequences with GenBank accession number *AAF20481*.*1*_GB_ and *AAF20475*.*1*_GB_ have both 110 amino acids and differ at two positions (91 and 99). Despite this, they share the same UniProt **CRC-64** checksum. The **SHA-1**-based SEGUID v2 checksums differ.

While collisions are unavoidable for all functions producing finite-length checksums, longer checksums are less likely to generate collisions. However, long checksums are less readable and also more expensive to generate, store, and compare. The SEquence Globally Unique IDentifier (SEGUID) checksum (Babnigg and Giometti, 2006) was introduced in order to provide a stable, unifying key for the same sequence in different databases facilitating linking protein sequences across databases. SEGUID is based on the **SHA-1** (Secure Hash Algorithm 1) algorithm (FIPS PUB 180-4, 2015), which produces a 160-bit hash, and is therefore less sensitive to collisions compared with the 64-bit **CRC-64** hash. With **SHA-1**, sequence collisions are very unlikely. For example, the two proteins with clashing **CRC-64** checksums (Figure 1) have distinct SEGUID checksums.

More importantly, no current checksum algorithm is directly useful for DNA, since the topology (linear vs circular) of DNA is most often different from that of proteins. Double-stranded DNA (dsDNA) consists of two complementary, anti-parallel strands. Usually, only one is stored for DNA, as the second strand is the complement of the stored sequence. However, both strands are equally valid representations of the DNA molecule, but give different checksum values, except for the special case where the sequence is a palindrome and the strands are therefore identical. DNA may be circular, e.g. circular single-stranded DNA (ssDNA) or circular dsDNA. For these cases, there are multiple representations for the same circular sequence, and there is no well-defined canonical representation for which the checksum should be calculated.

In this article, we propose SEGUID v2, which is an extension to the original SEGUID. Unlike existing algorithms, SEGUID v2 produces checksums for linear, circular, single- and double-stranded sequences of different types, including DNA, RNA, and amino-acid sequences in order to integrate synthetic biology resources and identify redundancies in existing databases.

We believe that a stable checksum that describes any biological sequence, regardless of topology or strandedness (single vs double strand), would streamline the exchange of information and biological materials within the synthetic biology community.

## 2 Methods

Defining a checksum for diverse sequence types involves three main challenges: (i) identify a unique representation among one or more equivalent representations, and convert it to a linear sequence representation, (ii) choose a hash function to distill the unique representation to a large-enough bit sequence, and (iii) compactly encode the bit sequence to a checksum using a text-friendly encoding alphabet.

Below, we explain how we define each of these steps for the proposed SEGUID v2 checksum, and what objectives we target when there are multiple options available.

### 2.1 Unique representation of sequence data

#### 2.1.1 Linear single-stranded sequences

A *linear single-stranded sequence* is a single, directional sequence with one or more symbols, where *directional* means it has a unique start and end (“left to right”). The sequence may be amino acids, DNA, RNA, or any sequence of symbols from a well-defined, case-sensitive alphabet. Examples of DNA sequences are `C, AT,` and `GATTACA`. Examples of RNA sequences are `C, AU,` and `AGA`. Examples of amino-acid sequences (“proteins”) are `A, QSA,` and `QSALTQPASV`.

A linear single-stranded sequence is always unique, which makes it natural to use the sequence as-is to calculate the checksum. This case was already covered by the original SEGUID algorithm.

#### 2.1.2 Linear double-stranded sequences

*Linear double-stranded sequences* comprise two sequences that share a region where the two sequences are complementary to each other. For double-stranded DNA, the 5′ → 3′ “upper” strand is often referred to as the “Watson” strand and the complementary “lower” as the “Crick” strand. We use the Watson and Crick strands separated by the symbol ↩, where both Watson and Crick are 5′ → 3′ sequences, to represent a double-stranded sequence. Using this notation, the double-stranded sequence `GATTACA` ↩ `TGTAATC` reflects the graphical representation in Figure 2.

**Figure 2:**
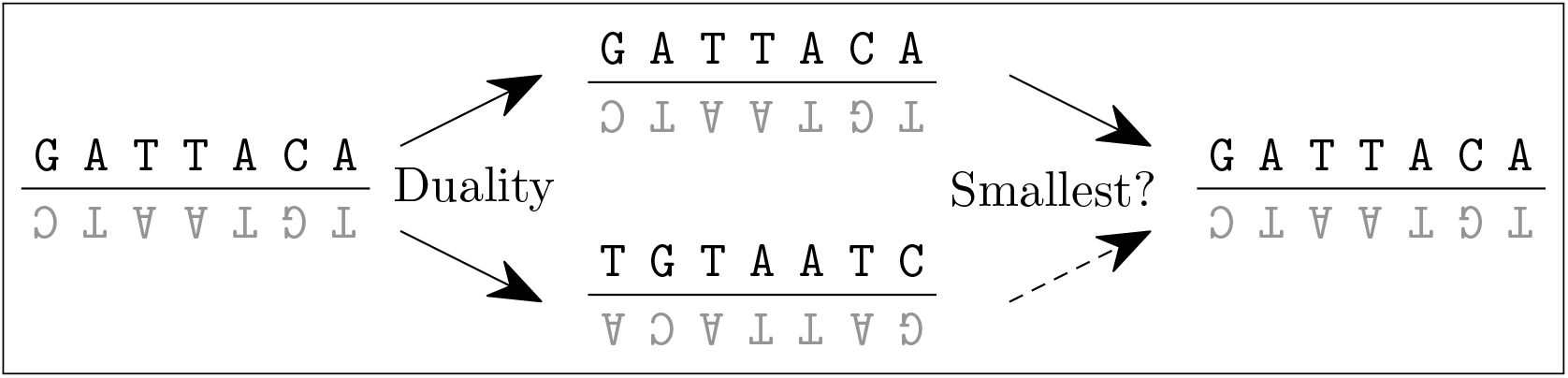
The SEGUID v2 steps for identifying the canonical representation of a blunt-ended linear, double-stranded DNA sequence. The input sequence `GATTACA` ←`TGTAATC` (left and top middle) has a dual, complementary representation `TGTAATC` ←`GATTACA` (bottom middle). The lexicographically smallest of these two is `GATTACA` ←`TGTAATC` (right). It is from this linear representation that the SEGUID v2 checksum is calculated. Legend: Following IUPAC-IUB Commission on Biochemical Nomenclature (1970), we choose to rotate the symbols on the complementary strand in the figure such that the left-to-right, 3′ → 5′ representation holds when rotating the figure upside-down.

Two types of such sequences must be considered - (i) blunt-ended, and (ii) staggered, linear double-stranded sequences - which are discussed next.

##### Blunt-ended, linear double-stranded sequences

In its basic form, the two sequences are of the same length and fully overlapping, i.e. the shared region spans their full length. We refer to this type of sequences as *blunt-ended* (short blunt), or non-staggered, double-stranded sequences. For example, `GATTACA` ↩`TGTAATC` is a blunt, double-stranded sequence with complementary strands `GATTACA` and `TGTAATC`. Importantly, `GATTACA` ↩`TGTAATC` and `TGTAATC` ↩`GATTACA` are dual representations of the same linear double-stranded DNA sequence (Figure 2). In other words, *without further constraints*, there is no unique representation of it. To address this *uniqueness problem*, we define the unique representation of the two possibilities as the one that is lexicographically smaller^5^ than the other. This is possible for all well-defined alphabets. For example, consider the DNA alphabet `A`, `C`, `G`, `T`, ordered such that `A` < `C` < `G` < `T`. This allows us to compare any two sequences lexicographically, e.g. AT < TA. For the above example, we have that `GATTACA` ↩`TGTAATC` < `TGTAATC` ↩`GATTACA`, and any checksums will be calculated based on the first of these two representations (Figure 2).

##### Staggered, linear double-stranded sequences

Another form of double-stranded sequences is where the two sequences do not fully overlap. We refer to this type of sequences as *staggered* double-stranded sequences, e.g.`-ATTACA` ↩`TGTAATC`, `--TTACA` ↩`TGTAATC`, and `-ATTACA` ↩`-GTAATC`. Here we use the minus symbol (-) to indicate that there is a symbol on the opposite strand but not on the current strand. As for blunt double-stranded sequences, staggered ones have two possible representations for the same sequence, e.g. `-GTAATC` ↩`--TTACA` and `--TTACA` ↩`-GTAATC` refer to the same sequence. As before, we define the unique representation to be the lexicographically smallest of the two, where we define the minus symbol (-) to be smaller than any other symbol, e.g. with the DNA alphabet we have that - < `A` < `C` < `G` < T. With this, we can identify `--TTACA`↩`-GTAATC` as the minimal lexicographic representation, because `--TTACA`↩`-GTAATC` < `-GTAATC`↩`--TTACA` (Figure 3). It is from this canonical representation that the hash is calculated.

**Figure 3:**
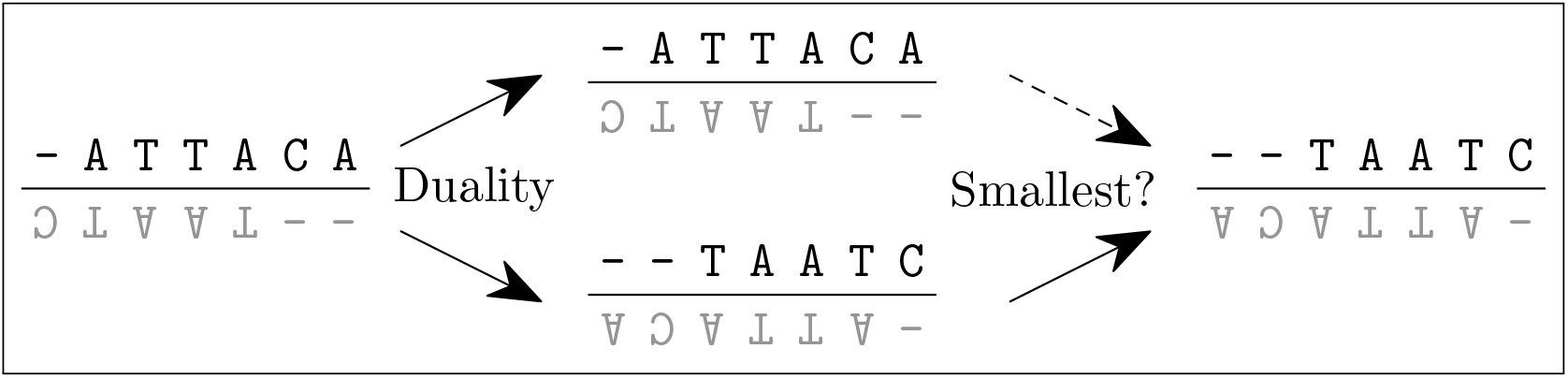
The SEGUID v2 steps for identifying the canonical representation of a staggered, linear, double-stranded DNA sequence. The input sequence `-ATTACA` ↩`--TAATC` (left and top middle) has a dual, complementary representation `--TAATC` ↩`-ATTACA` (bottom middle). Because the minus symbol (-) is lexicographically smaller than the other symbols, the smallest of these two linear representations is `--TTACA` ↩`-GTAATC` (right). It is from this linear representation that the SEGUID v2 checksum is calculated.

#### 2.1.3 Circular single-stranded sequences

A *circular single-stranded sequence* is a single, directional sequence with one or more symbols without a beginning or end. The sequence may comprise symbols from any well-defined alphabet. Any linear single-stranded sequence can be made a circular single-stranded sequence, by combining the two ends, and vice versa. Examples of circular DNA sequences are `C`⟳, `AT`⟳, `TA`⟳, and `GATTACA`⟳. Here we use the symbol ⟳ to indicate that the two ends are combined. A naturally occurring circular, single-stranded DNA sequence is the 6,407-nucleotide M13 bacteriophage with sequence AACGCTACTA…GGATGTT⟳ (*NC_003287*.*2*_GB_ and *V00604*.*2*_GB_). Circular RNA (circRNA) is naturally occurring circular RNA formed from mRNA through a process called back-splicing (Jeck et al., 2013), where a downstream splice donor site is joined to an upstream splice acceptor site. The *MW729338*.*1*_GB_ represents an 840-nucleotide circRNA from *Culex pipiens* (northern house mosquito). Examples of circular proteins are `A`⟳, `QSA`⟳, and `QSALTQPASV`⟳. A naturally occurring circular single-stranded amino-acid sequence is the 21 amino-acid long *E. coli* peptide Microcin J25 (*AAD28494*.*1*_GB_ at positions 38–58) with sequence VGIGTPISFYGGGAGHVPEY⟳. This circular peptide is also referred to as VGIGTPISFYGGGAGHVPEYF⟳, e.g. Blond et al. (1999) and Trabi and Craik (2002).

Because there is no beginning and end of a circular sequence, there is also no unique representation of a circular sequence *without further constraints*. Both `AT`⟳ and `TA`⟳ represent the same circular sequence. Similarly, `GATTACA`⟳, `ATTACAG`⟳, `TTACAGA`⟳, `TACAGAT`⟳, `ACAGATT`⟳, `CAGATTA`⟳, and `AGATTAC`⟳ are all the same circular sequence. In general, there are *n* different representations of the same circular single-stranded sequence of length *n*.

The non-uniqueness of these representations complicates how to store a circular sequence in databases, how to communicate it electronically, and how to present it in text. Similarly, if we naively would calculate the original SEGUID checksum for the above representations, we would get different checksums despite them representing the same circular sequence.

To address the above *uniqueness problem*, we need to identify a single unique representation among all possible representations part of the same *equivalent class*. Analogously to how we addressed the uniqueness problem for double-stranded sequences, a natural choice is to define the unique representation to be *the sequence with a rotation that has the smallest lexicographical value*. For example, consider the above circular sequence `GATTACA`⟳ with seven equivalent representations. We can order the seven possible rotations as `ACAGATT` < `AGATTAC` < `ATTACAG` < `CAGATTA` < `GATTACA`< `TACAGAT` < `TTACAGA`. The *minimal lexicographical rotation* of `GATTACA`⟳ is therefore `ACAGATT` (Figure 4). The minimal rotation can be found in linear time (Pierre Duval, 1983).

**Figure 4:**
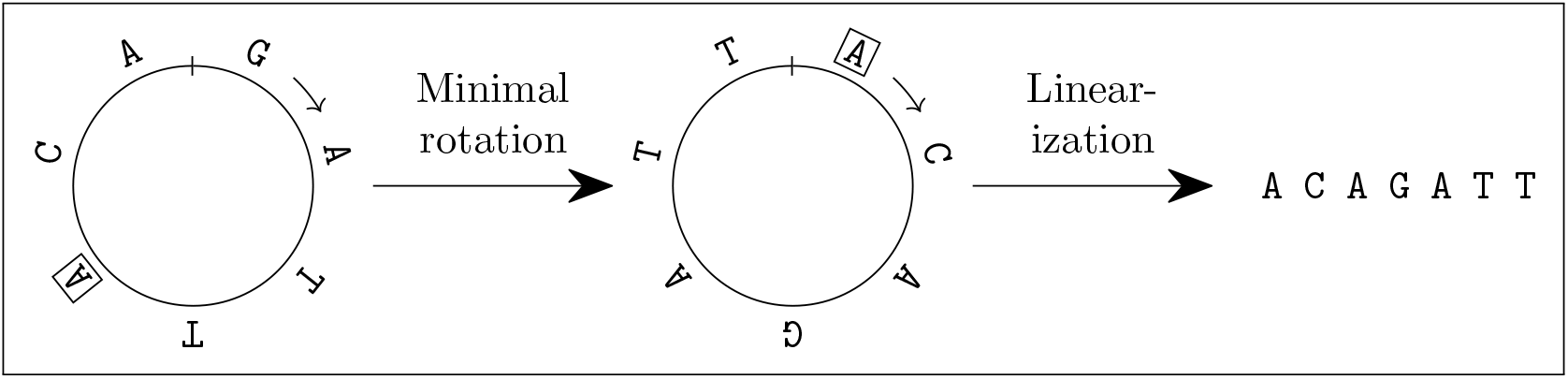
The SEGUID v2 steps for identifying the canonical representation of a circular, single-stranded DNA sequence. The input sequence `GATTACA`⟳ (left) is one of several equivalent, non-normalized representations of the same circular sequence. Among all seven possible rotations, there is a minimal rotation, which for this sequence is `ACAGATT`⟳ (middle). The linear representation of the minimal rotation is `ACAGATT` (right). It is from this linear representation that the SEGUID v2 checksum is calculated.

#### 2.1.4 Circular double-stranded sequences

A *circular double-stranded sequence* is similar to a *circular single-stranded sequence*, but with a second complementary strand. Examples of circular double-stranded DNA sequences are `C` ↩ `G`⟳, `AT` ↩ `AT`⟳, and TATGCCAA ↩TTGGCATA⟳.

A real-world example is the 2,686-basepair circular double-stranded DNA pUC19 cloning vector TCGC…CGTC↩ GACG…GCGA⟳ (*M77789*.*2*_GB_ and *L09137*.*2*_GB_).

Similarly to the single-stranded cases, there are multiple representations for circular double-stranded sequences. For example, the eight rotational variants TATGCCAA ↩TTGGCATA⟳, ATGCCAAT ↩ATTGGCAT⟳, TGCCAATA ↩TATTGGCA⟳,…, ATATGCCA ↩TGGCATAT⟳ represent the same eight-basepair circular dsDNA, but so does the eight complementary counterparts TTGGCATA ↩TATGCCAA⟳, ATTGGCAT ↩ATGCCAAT⟳, TATTGGCA ↩TGCCAATA⟳, …, TGGCATAT ↩ATATGCCA⟳. In general, there are 2 · *n* different representations of the same circular double-stranded sequence of length *n*. We define the unique representation as *the smallest of the minimal lexicographical rotations of the two strands*. For instance, AATATGCC↩GGCATATT⟳ is the minimal for one of the original representation, and ATATTGGC↩GCCAATAT⟳ for the complementary duality. Between these two, AATATGCC↩GGCATATT⟳ is the lexicographically small-est, because AATATGCC ↩GGCATATT⟳ < ATATTGGC ↩GCCAATAT⟳. Thus, the canonical representation of the circular double-stranded sequence TATGCCAA ↩TTGGCATA⟳ is AATATGCC ↩GGCATATT⟳ (Figure 5). Note, it is sufficient to compare only AATATGCC < ATATTGGC to identify AATATGCC ↩GGCATATT⟳ to be the lexicographically minimal representation. This is true also for non-bijective complementary alphabets.

**Figure 5:**
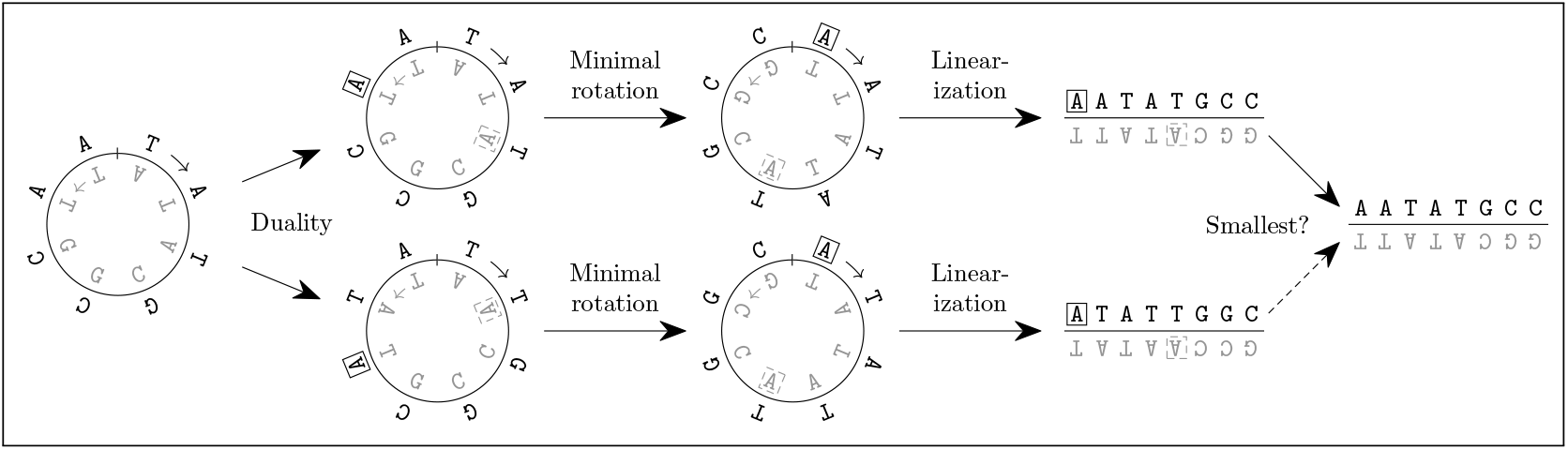
The SEGUID v2 steps for identifying the canonical representation of a circular, double-stranded DNA sequence. The input sequence TATGCCAA ↩TTGGCATA⟳ (left, and top left) is one of several equivalent, non-normalized representations of the same circular sequence. This double-stranded sequence has the dual complementary representation TTGGCATA ↩TATGCCAA⟳ (bottom left). The minimal rotations for these dual representations are AATATGCC ↩GGCATATT⟳ (top middle) and ATATTGGC ↩GCCAATAT⟳ (bottom middle). The smallest of the two linear minimal-rotated representations (top right and bottom right), is AATATGCC ↩GGCATATT (right). It is from this linear representation that the SEGUID v2 checksum is calculated.

### 2.2 Hash function (SHA-1)

SEGUID v2 supports linear and circular, single- and double-stranded sequences. As shown above, circular sequences are transformed to unique, linear representations based on minimal rotations and lexicographic ordering. Thus, regardless of input sequence, the hash is always calculated on a linear representation. To apply the hash function to these representations, we need to coerce them to single strings^6^. There are two types of linear representations we have to consider; the single-stranded one and the double-stranded one. A linear *single*-stranded sequence can be coerced to a string as-is, e.g. the sequence `GATTACA` becomes the string “`GATTACA`”. For a linear *double*-stranded sequence, we chose to coerce it to a string by concatenating the 5′ → 3′ Watson sequence and the 5′ → 3′ Crick sequence with a semicolon (;). For example, for the blunt, linear dsDNA sequence `GATTACA` ↩`TGTAATC`, the hashed string is “`GATTACA`;`TGTAATC`”. Similarly, for the staggered, linear ds-DNA sequence `--TTACA` ↩`-GTAATC` the hashed string is “`--TTACA`;`-GTAATC`”. The original SEGUID method uses **SHA-1** to generate a 160-bit hash from the input sequence. As argued in Babnigg and Giometti (2006), **SHA-1** is less sensitive to collisions compared to **CRC-64** and sequence checksum collisions are very unlikely when using **SHA-1**. Another popular hash function is the Message Digest Algorithm 5 (**MD5**) (Rivest, 1992), which produces a 128-bit hash. The UniProt database uses the **MD5** checksum. Although we are not aware of any collisions for biological data, it has been shown (Wang and Yu, 2005) that **MD5** collisions can be created artificially. Although we have not yet observed a **SHA-1** collision, one could argue that we should “upgrade” to a **SHA-2** algorithm to further lower the risk for collisions. On the other hand, that comes with a cost of producing longer checksums. For example, the popular and widely available **SHA-256** algorithm produces a 256-bit long hash, which is 60% longer than a **SHA-1** hash, which in turn would lengthen the final checksum the same amount. There are downsides to having long checksums. For instance, they may be too long to be used for metadata headers in some sequence file formats, they risk being line-wrapped in communications such as email threads, and they add to the storage size for very-large lookup tables. Moreover, some file systems have a limit on how long a canonical pathname may be, e.g. on Microsoft Windows the default pathname-length limit is 256 characters. A very long checksum in a directory or file name, in an already deeply nested directory structure, may exceed this limit. An alternative to **SHA-256** is **SHA-224**, which results in “only” a 40% longer checksum and might therefore be a hash-algorithm candidate. Unfortunately, implementations of **SHA-224** are less common across programming languages than **SHA-1, SHA-256**, and **SHA-512**. Yet another alternative is **SHA-512/224**, which produces a 224-bit truncated hash based on the **SHA-512** algorithm (FIPS PUB 180-4, 2015). The SHA specification supports **SHA-512/t** for other values of **t**, e.g. **SHA-512/160** would produce a hash of the same length as **SHA-1**. Readily available implementations of these truncated alternatives are even less common than **SHA-224**. An alternative approach to **SHA-512/t** was proposed by Hart and Prlić (2020), where they discard the last 320 bits of the 512-bit **SHA-512** hash to produce a 192-bit hash. We contemplated this approach, but we decided against it as we do not know if it is statistically sound or valid for our needs.

Lastly, Stevens et al. (2017) and Leurent and Peyrin (2020) demonstrated that it is feasible to artificially construct **SHA-1** clashes on modern computers. Because of this, **SHA-1** is no longer considered safe from a cryptography point of view. For instance, the U.S. National Institute of Standards and Technology (NIST) “*recommends that federal agencies transition away from SHA-1 for all applications as soon as possible. Federal agencies should use SHA-2 or SHA-3 as an alternative to SHA-1*” (Computer Security Resource Center (CSRC), 2022). The **SHA-2** and **SHA-3** family of hash functions produce longer hashes, making it harder to construct collisions. That said, sequence checksums are used for efficient storage, communication, and lookup of sequences - not to secure them cryptographically. Arguing that the checksum of a full sequence can guarantee that the full sequence has not been compromised is problematic and risky. To achieve that, cryptographical protocols based on, for instance, public-and-private keys should be used, which is a problem beyond sequence checksums. We find no reason to incorporate such features in SEGUID v2, because there already exist well-established, readily available protocols for encrypted communication and storage.

For these reasons, we choose to use **SHA-1** also for SEGUID v2.

### 2.3 Encoding (Base64url)

A hash, by itself, is sufficient to uniquely identify a sequence. However, hashes, such as **SHA-1**, are very large integers, larger than can easily be represented in common programming languages, which makes them complicated to coerce as-is to and from textual representations. Encoding is a technique to turn an internal binary representation to a string, preferably in a compact way. Different encoding algorithms have different “compression” ratios (Table 1).

**Table 1:**
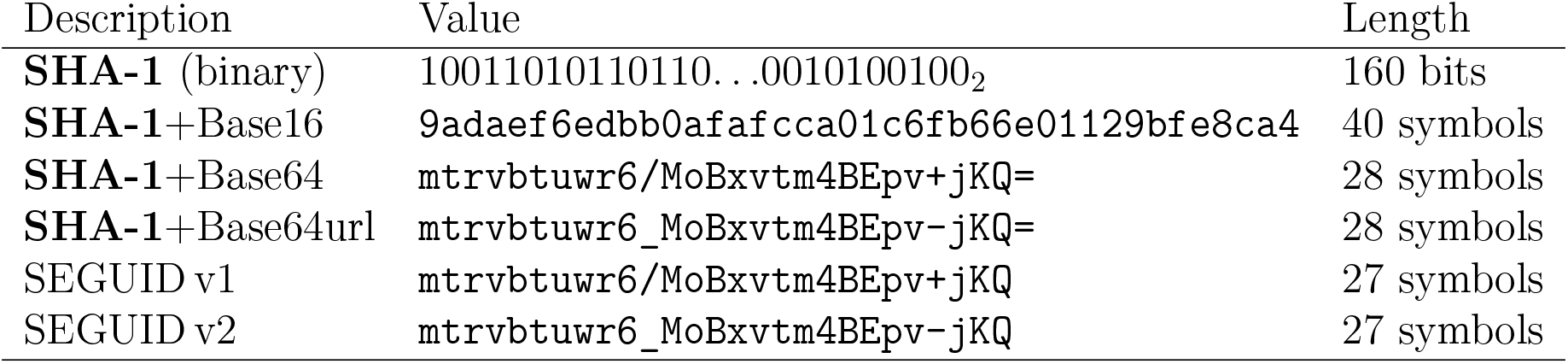
Examples of different encodings of a SHA-1 hash. The **SHA-1** hash was generated based on the sequence string `ACAGATT`. The **Base64** specification requires that the output comprise a multiple of four symbols, and if not, the output should be padded with equal signs (=) at the end. Because of this, a **Base64** encoding of a **SHA-1** hash will always have a single, non-informative padding symbol at the end. This padding symbol is excluded in the SEGUID checksums.

For instance, the **SHA-1** hash, which is a 160-bit integer, or equivalently a 20-byte integer, could be encoded to a string of length 160 comprising symbols zero (0) and one (1). Assuming an 8-bit ASCII string, this encoding is 800% larger than the internal 20 bytes. That would be an inefficient way to represent the hash. A more compact, and commonly used encoding is the hexadecimal representation, which comprise a 16-character alphabet (0-9 and `a-f`). In this representation, a **SHA-1** hash is of length 40 characters; one for every four (log_2_ 16) bits, or two for each byte. To achieve a more compact checksum, a larger set of symbols is needed. The **Base64** encoding (Josefsson, 2006) uses a 64-character alphabet (`A-Z`, `a-z`, `0-9`, +, and /). With this, each character can encode six (log_2_ 64) bits, i.e. a 160 bits can be represented by ⌈160*/*6⌉ = 27 characters.^7^

The original SEGUID method uses **Base64**, modulo the mandatory, non-informative padding symbol, to encode the **SHA-1** hash, resulting in a 27-character checksum. This was chosen to get a compact representation of the hash, while still being text-friendly. One problem with a **Base64**-based checksum is that it cannot be used as part of a filename, because of the forward slash (/), which is used to represent directories on most file systems. For a similar reason, it cannot be used as-is as part of a Uniform Resource Locator (URL), where symbols / and + have special meaning. In order to use a SEGUID checksum in a URL, these symbols need to be re-encoded. To address these needs, a filename- and URL-safe variant of **Base64** is the **Base64url** encoding (Josefsson, 2006). It substitutes forward slashes (/) and plus symbols (+) with underscores (_) and minus symbols (-). The **Base64url** algorithm is linear (*O*(*n*)) in the number of bits, which is always 160 bits when using **SHA-1**.

In contrast to the original SEGUID, for SEGUID v2, we choose to use the **Base64url** encoding. For example, the linear single-stranded DNA sequence `ACAGATT` yields the SEGUID v1 checksum `mtrvbtuwr6/MoBxvtm4BEpv+jKQ`, while the SEGUID v2 checksum is `mtrvbtuwr6_MoBxvtm4BEpv-jKQ` (Table 1).

### 2.4 SEGUID v2 checksums have a prefix

In order to preserve what type of information is encoded in the checksum, SEGUID v2 checksums are by default prefixed with either `lsseguid, csseguid, ldseguid`, or `cdseguid`, followed by an equal sign (=). For example, `lsseguid=IQiZThf2zKn_I1KtqStlEdsHYDQ` indicates a SEGUID v2 checksum for a linear single-strand sequence.

The prefix helps to distinguish SEGUID v2 checksums from other types of 27-character checksum strings. It also conveys the type of sequence the checksum represents. We have found that the latter helps users to detect mistakes sooner in case they selected the incorrect sequence type for SEGUID v2.

Another advantage of the prefix is that it eliminates the risk of getting checksums with all digits, which some software tools might parse as numeric values. For example, Microsoft Excel and LibreOffice Calc will coerce such all-digit strings to numerics, but with loss of precision. If such a numeric value is coerced back to a string, it will comprise the first 15-16 digits followed by all zeros. We observe the same problem with higher-level programming functions (e.g. in R and Python) that read data from file risks reading all-digit columns as loss-of-precision numeric values or overflowed integer values.

### 2.5 SEGUID Short ID

For practical purposes, SEGUID v2 defines a *short form* of its checksum. Specifically, a *SEGUID Short ID* comprises the first six (6) characters of the checksum without the prefix. For example, the Short ID of the SEGUID v2 checksum `lsseguid=IQiZThf2zKn_I1KtqStlEdsHYDQ` is `IQiZTh`.

While we designed the SEGUID v2 checksum to be text-friendly, we designed the Short ID to be human-friendly. The *SEGUID Short ID* provides a compact way to reference a specific sequence. The short length makes it easy to communicate in both written and spoken form, and they allow for quick manual comparisons. It is also feasible to memorize and recognize, especially in projects that have a few sequences that are commonly used.

The risk for collisions among Short IDs is still relatively small, especially for any given project working with a limited set of sequences. Theoretically, a six-character Short ID can encode 2^36^ = 1, 073, 741, 824 different sequences^8^.

### 2.6 Supported sequence types and alphabets

The SEGUID v2 algorithm is designed to support DNA, RNA, and proteins, but also sequences with a custom alphabet (Table 2). Formally, the input sequence must not include symbols outside of the declared *case-sensitive* alphabet. By requiring case sensitivity, we allow for a richer set of custom alphabets, but we also avoid the risk of introducing non-deterministic behavior from, for instance, automatically coercing lower-case letters to upper-case ones. In addition, we limit alphabets to digits, and upper-, and lower-case letters. This is to assure a well-defined lexicographic ordering of symbols (per the 7-bit ASCII character set). Importantly, there are *reserved symbols* that must never be part of the alphabet. Specifically, the dash symbol (-), the semicolon (;), and the newline (`\n`) are reserved symbols.

**Table 2:**
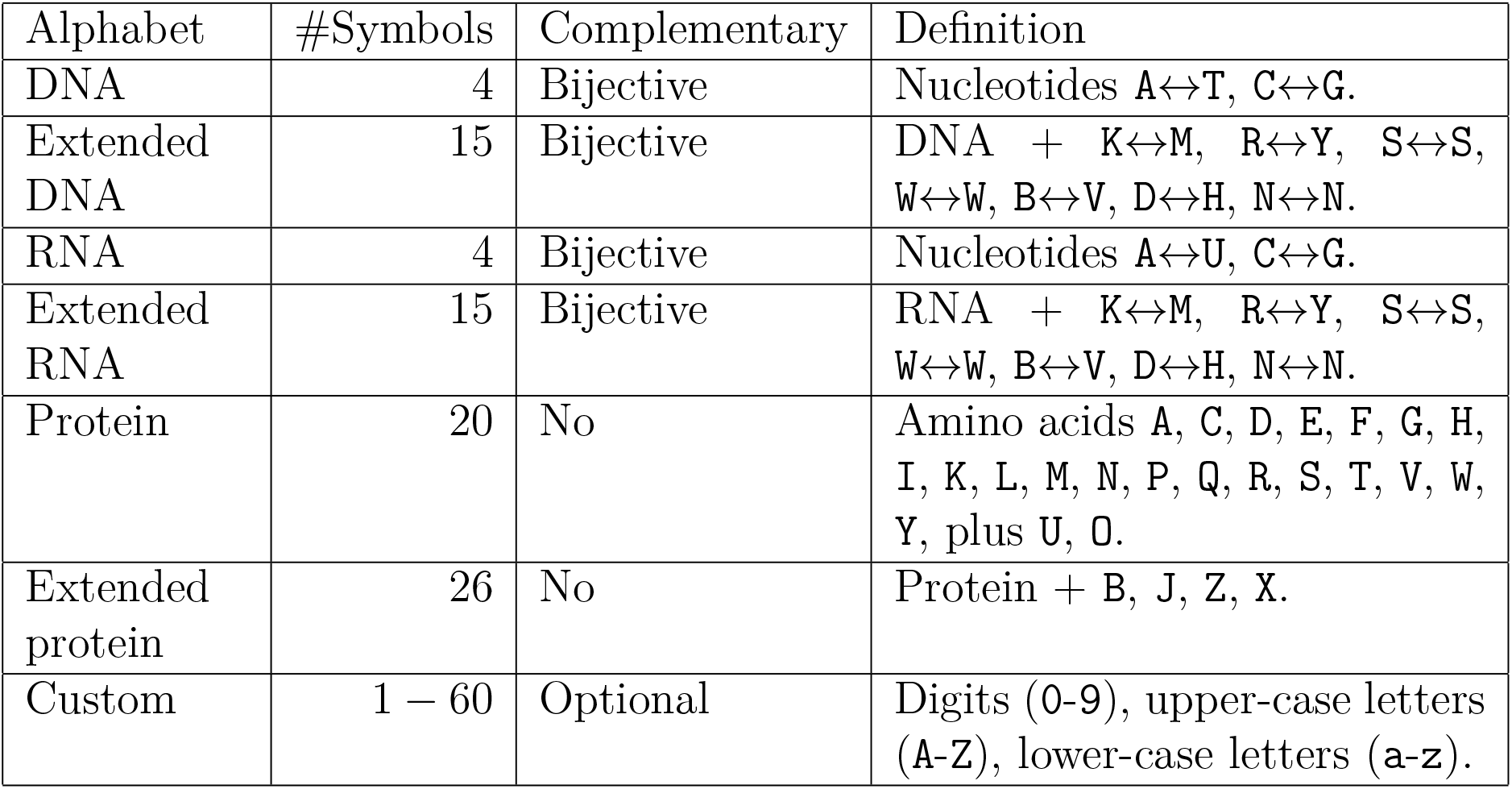
Alphabets supported by SEGUID v2. SEGUID v2 has built-in support for DNA, RNA, and proteins. These alphabets comprise upper-case letters only. IUPAC is a standard to represent DNA, RNA, and amino-acid sequences with an “extended” set of symbols (IUPAC-IUB Commission on Biochemical Nomenclature, 1970, 1984). Custom alphabets, complementary or not, are also supported. The concept of complementarity does not apply to proteins. Proteins are effectively single-stranded sequences, meaning only the single-stranded SEGUID v2 methods apply to them. Amino-acid symbols `J`, `O`, and `U` are not part of the IUPAC standard.

## 3 Results

SEGUID v2 is designed to be robust against mistakes in order to prevent incorrect checksums from propagating in the scientific process. For instance, part of the requirements is that the implementation should validate the correctness of the input sequence, e.g. all symbols are part of the alphabet and double-stranded sequences are correctly complementary. At the same time as being strict, SEGUID v2 is designed to be flexible such that it can support existing and future sequences not already anticipated. For example, in Viner et al. (2024), the authors define an expanded epigenetic DNA alphabet (Table 3) that encodes also known modifications to cytosine (C). Using this alphabet, methylation motifs can be represented as, for instance, `TTGmGCAA`, `AT2hAAAT`, and `TGCm1m1`. Their complementary counterparts are `TTGC1CAA`, `ATTT2hAT`, and `m1m1GCA`. They also define an “extended” version of this alphabet, with additional complementary symbol pairs. Both of these alphabets can be used with SEGUID v2 by defining them as custom alphabets.

**Table 3:**
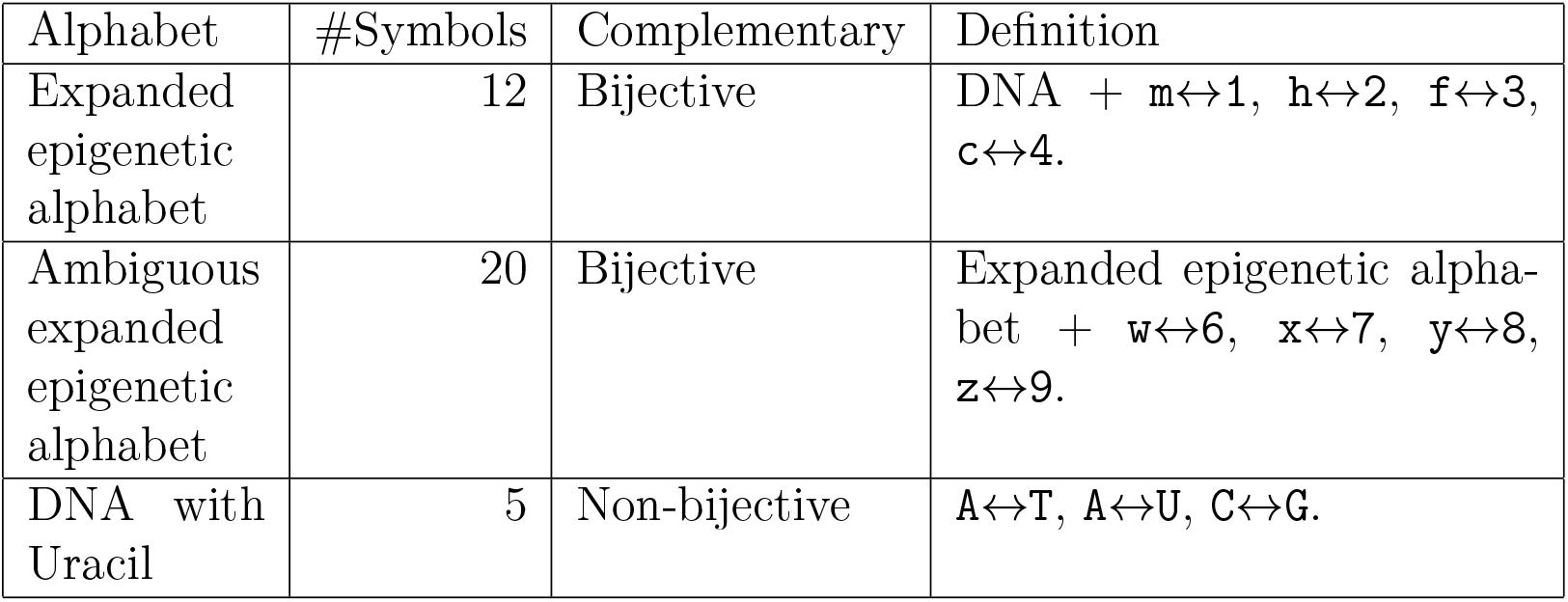
Custom alphabets with SEGUID v2. (i) The expanded epigenetic alphabet, defined in Table 1 of Viner et al. (2024), includes known modifications to cytosine (`C`) and symbols for each guanine (`G`) complementary to a modified nucleobase. (ii) The “ambiguous” version of this alphabet, defined in Table 2 of Viner et al. (2024), addresses the case when the knowledge on some of the nucleotides is incomplete. (iii) Uracil (`U`) may occur naturally in DNA, which in case adenine (`A`) is complementary to either thymine (`T`) or uracil (`U`). This can be represented by a non-bijective complementary alphabet.

Another example arises from the fact that uracil (U) may occur naturally in DNA, e.g. phage DNA from dUTPase, uracil-DNA glycosylase (*dut ung*) mutant hosts (Wang and Mosbaugh, 1988), as an apoptosis inducer during *Holometabola* development (Muha et al., 2012), and as a means for somatic hypermutation resulting from U:G mismatch repair during vertebrate antibody development (Muramatsu et al., 2000). See Figure 6 for an illustration.

**Figure 6:**
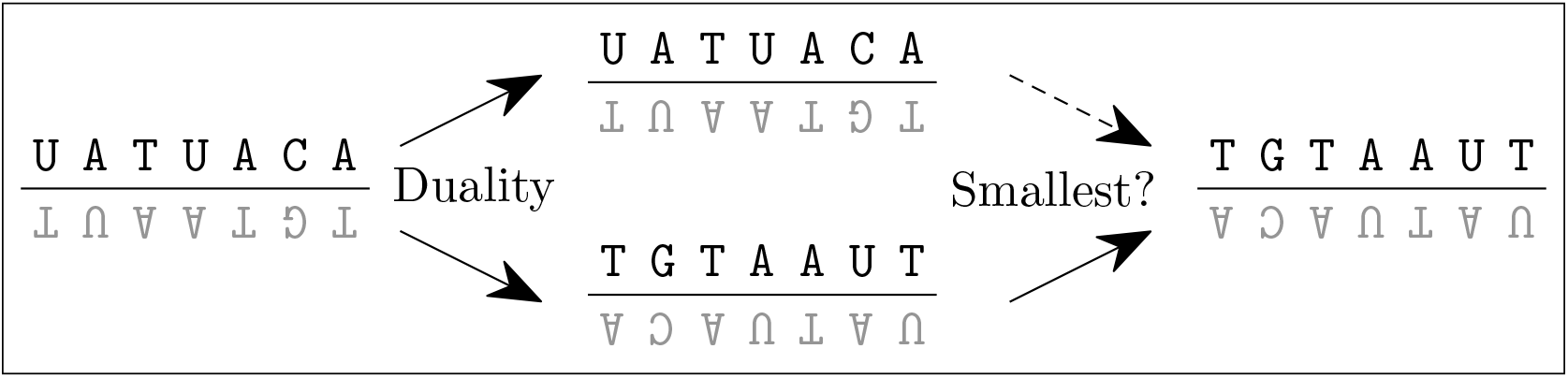
The SEGUID v2 algorithm on double-stranded sequence of DNA with uracil. The SEGUID v2 algorithm supports non-bijective complementary alphabets such as DNA with uracil (U). The input sequence `UATUACA` ↩ `TGTAAUT` (left and top middle) comprise basepairs `A`↔ `T` and `A`↔ `U`. It has a dual, complementary representation `TGTAAUT` ↩ `UATUACA` (bottom middle), which is also the lexicographically smallest (right).

This also occurs in the USER™ Uracil excision cloning protocol (Geu-Flores et al., 2007). In these cases, adenine (A) may bind to either thymine (T) or uracil (U), which can be described by a *non-bijective* complementary alphabet (Table 3).

## 4 Implementation and availability

### 4.1 Minimal, core software implementations

The SEGUID v2 algorithm, together with the original SEGUID algorithm, is implemented and distributed as libre, open-source software under the MIT license. We provide small, self-contained implementations in JavaScript (ECMA International, 2024), Python (van Rossum and de Boer, 1991), R (R Core Team, 2024), and Tcl (Ousterhout and Jones, 2009) that have a minimal set of dependencies. For JavaScript, Python and R, we provide packages named **seguid**, which are available on npm (node package manager), PyPi (Python Package Index), and CRAN (Comprehensive R Archive Network), respectively. These packages implement a standardized application protocol interface (API), e.g. `csseguid` (“`GATTACA`”). They also provide a command-line interface (CLI), e.g.

~~~
$ npx seguid --type=lsseguid <<< ↩GATTACA’
 lsseguid=tp2jzeCM2e3W4yxtrrx09CMKa_8
$ python -m seguid --type=cdseguid <<< ↩TATGCCAA;TTGGCATA’
 cdseguid=dUxN7YQyVInv3oDcvz8ByupL44A
$ Rscript -e seguid::seguid --type=csseguid <<< ↩GATTACA’
csseguid=mtrvbtuwr6_MoBxvtm4BEpv-jKQ
$ tclsh seguid --type=ldseguid <<< ↩TATGCCAA;TTGGCATA’
ldseguid=p88RYs41n0NTej4htM1fJAWI1ME
~~~

Documentation, examples, and online WebAssembly-based (Haas et al., 2017) demos are available on the SEGUID website (https://www.seguid.org). Support for additional programming languages is planned, specifically C++ and Rust, and will be made available via the SEGUID website.

### 4.2 Validation and testing

It is essential that any implementation of the SEGUID v2 checksum algorithm is correct and complies with the SEGUID v2 specifications as described here. For instance, we do not want incorrectly produced checksums to be used in production, appear in publications, or enter shared databases. We provide a rich set of unit and redundancy tests (see SEGUID website) to lower the risk for this to ever occur. These tests assert correctness, and identical behavior and results regardless of which programming language was used for the implementation. All new implementations must adhere to these tests in order to be considered SEGUID v2 compliant. These tests run on all releases and on a nightly basis. This test framework is designed to make it as simple as possible for a third-party contributor to run these tests. We use continuous-integration testing to validate all known implementations.

### 4.3 Higher-level implementations

The main purpose of the above **seguid** packages is to provide a minimal, language-agnostic API of the SEGUID v2 algorithm that can be imported in higher-level implementations. The scope of these packages will be kept at a minimum to avoid feature creep of both of the API and the SEGUID v2 algorithm. This way, the packages also serve as reference implementations and help lower the threshold for implementing the API in other languages.

For these reasons, we consider tasks like coercing a DNA sequence in lower-case letters to upper-case letters, trimming strings, and converting a sequence to its complementary counterpart to be out of scope. Such features have to be implemented by other tools and packages, e.g. Python packages **pydna** (Pereira et al., 2015) and **Biopython** (Cock et al., 2009; Bassi and Gonzalez, 2007). For example, **pydna** implements the SEGUID v2 methods for the Dseqrecord class, which holds double-stranded DNA. It also provides a rich set of utility functions for operating on DNA and RNA sequences. **Biopython** already implements the original SEGUID method and we anticipate it will also provide SEGUID v2 functionalities in a future release.

### 4.4 Graphical user interfaces (GUIs)

Some users are not comfortable with the provided command-line interfaces, or inexperienced with the supported programming languages. In order to make SEGUID v2 accessible to these users, it is important the method is also available through a Graphical User Interface (GUI). A readily available GUI will boost the adoption of SEGUID v2 to a broader audience as it also caters to users of varying expertise levels. Below are two GUIs that can be used for calculating SEGUID checksums.

The **SEGUID Calculator** (Figure 7) is a browser-based GUI for the SEGUID v2 algorithm available via the SEGUID website. It is implemented in JavaScript and allows for computations such that entered data never leaves the web browser or the local computer. This type of instant interactive feedback enhances understanding of how the checksum works.

**Figure 7:**
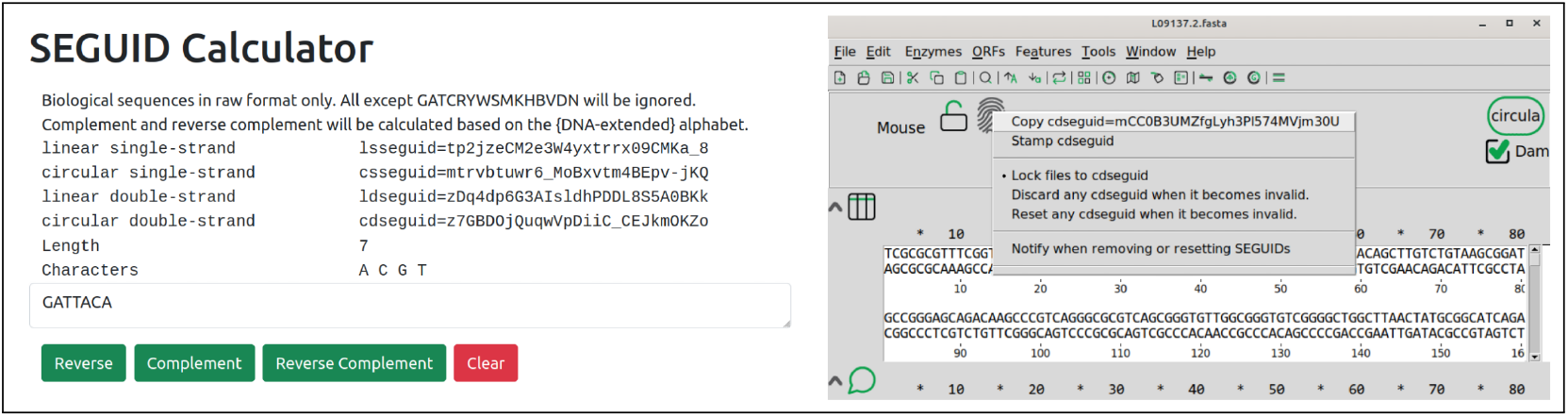
GUI implementations of SEGUID checksums. Two screenshots of GUI tools for calculating SEGUID v2 checksums. The **SEGUID Calculator** tool (left) calculates the different types of SEGUID v2 checksums based on the Watson sequence as it is being entered. This is a JavaScript-based implementation that fully runs within the web browser. The A plasmid Editor (**ApE**) tool (right) is specifically designed for double-stranded DNA. Because of this, it is sufficient to enter the Watson sequence, from which the Crick sequence is automatically inferred.

The popular, freely available **ApE** (A plasmid Editor) (Davis and Jorgensen, 2022) allows calculating and stamping dsDNA sequences with SEGUID v2 checksums (Figure 7). **ApE** is built on our Tcl implementation and available from the Jorgensen Lab website^9^.

## 5 Applications

SEGUID checksums can be used to map sequence metadata both within and across biological databases. For example, by indexing different types of identifiers based on SEGUID checksums, the SEGUID checksum can serve as the common denominator across heterogeneous databases. For example, this technique was used at the Argonne National Laboratory to build a real-time, interactive search engine for protein annotation data collected from multiple resources. For motivations similar to those of the Short ID, the search engine was designed to list matching candidates as soon as the user had typed three letters (Figure 8).

**Figure 8:**
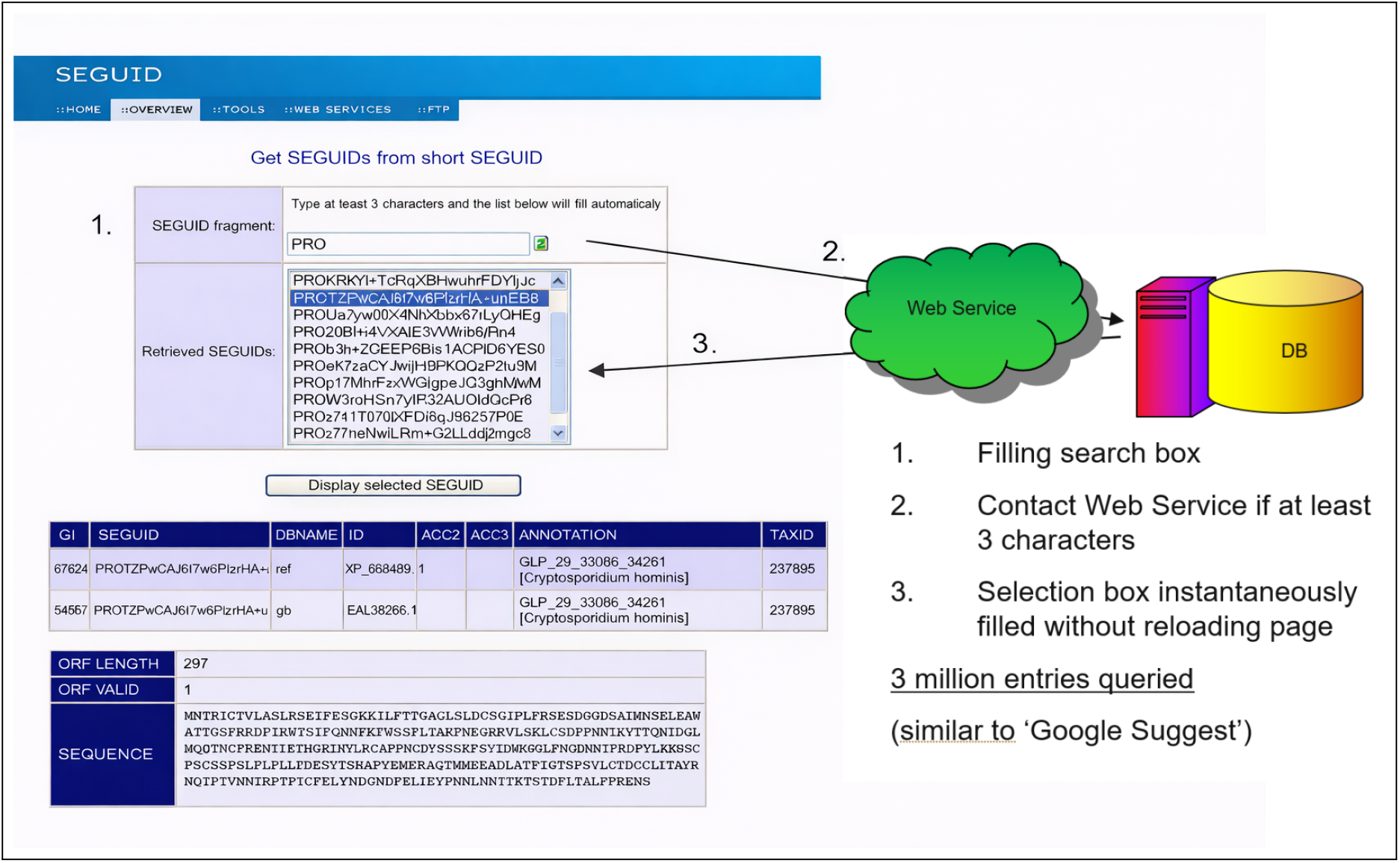
Type-ahead database search based on partial SEGUID checksums. Illustration of how partial SEGUID checksums can be used to autocomplete searches as you type. In this example, the user enters `PRO` and the Ajax-based web interface instantaneously returns all SEGUID checksums starting with those characters. Sequence data is pulled from multiple databases concurrently for entries matching the selected SEGUID checksum.

Analogously to how SEGUID checksums can be used for quick lookup of sequence data via database tools, they can be used for navigating and looking up sequence data hosted directly on the file system. For example, sequence-annotation data files of different kinds can be grouped in folders named as the corresponding SEGUID v2 checksums, or the individual files can comprise the checksum.

The **ApE** tool and the **pydna** package can stamp and save sequence files in GenBank format^10^ where the checksum is saved in the `COMMENT` section of the GenBank file (Figure 9). An existing checksum can be verified by each tool.

**Figure 9:**
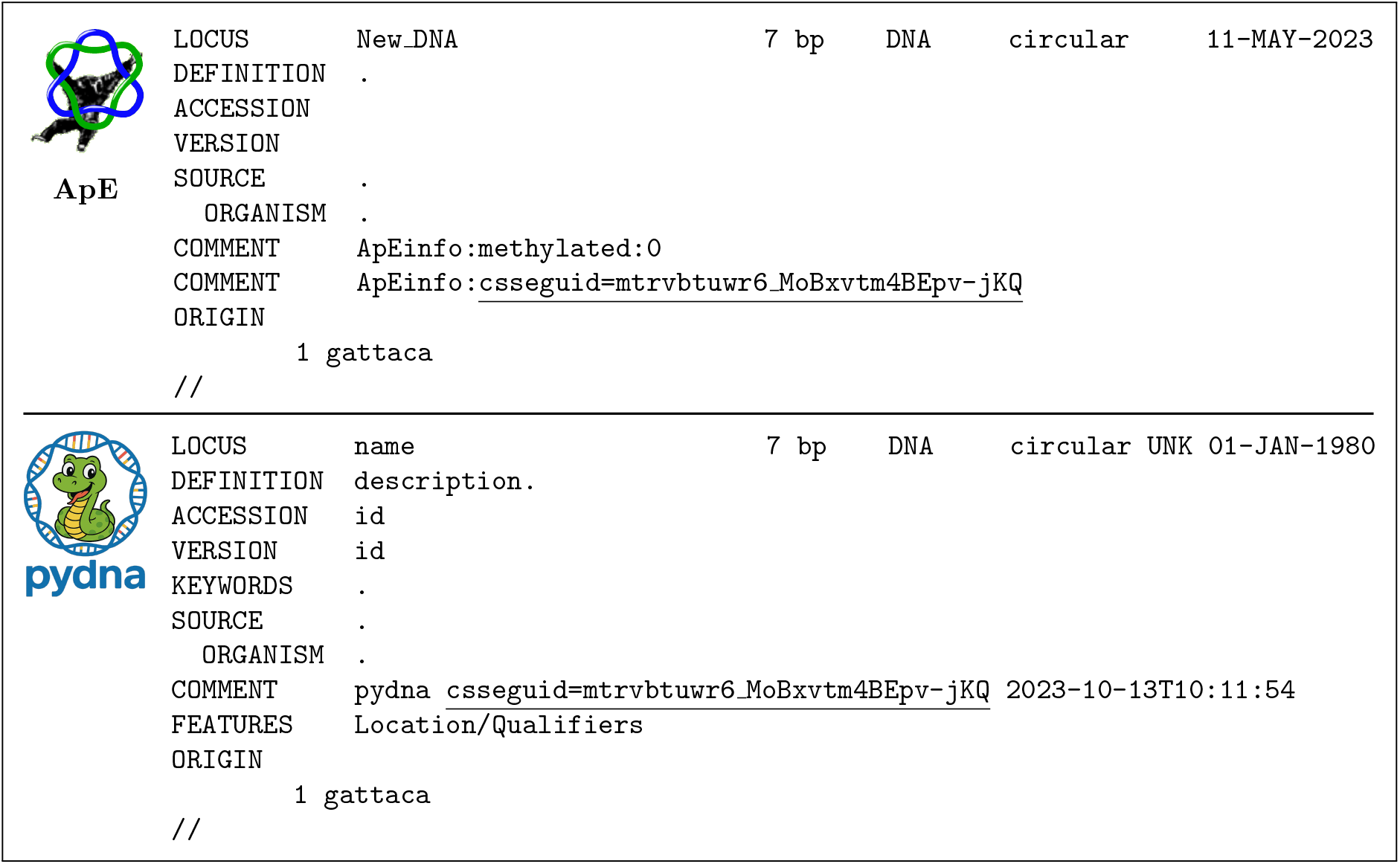
SEGUID checksums added to GenBank sequence files. The files were stamped with SEGUID checksum using the **ApE** tool (upper panel) or the **pydna** Python package (lower panel). In both cases, the checksum is added in the `COMMENT` field of the GenBank record. In addition, **pydna** appends an ISO-8601 timestamp after the checksum.

A common use of the SEGUID checksum is in undergraduate teaching of practical molecular biology in the Department of Biology at the University of Minho. An example problem is depicted in Figure 10. The students are given a task, typically to simulate a sub-cloning problem on their computers. Since this kind of problem has a unique correct answer, checksums for the correct solution can be calculated beforehand. The problem of assessing the students is simplified by the students’ access to a local or web-based SEGUID-capable software, e.g. **SEGUID Calculator**. Giving access to a partial checksum for the correct answer can help the students to rapidly determine whether their procedures are incorrect. We have found this approach practical to teach in-silico cloning to over 60 students at a time. Overall, precalculated SEGUID codes lower the number of mistakes filed and the efforts spent on reviewing homework assignments.

**Figure 10:**
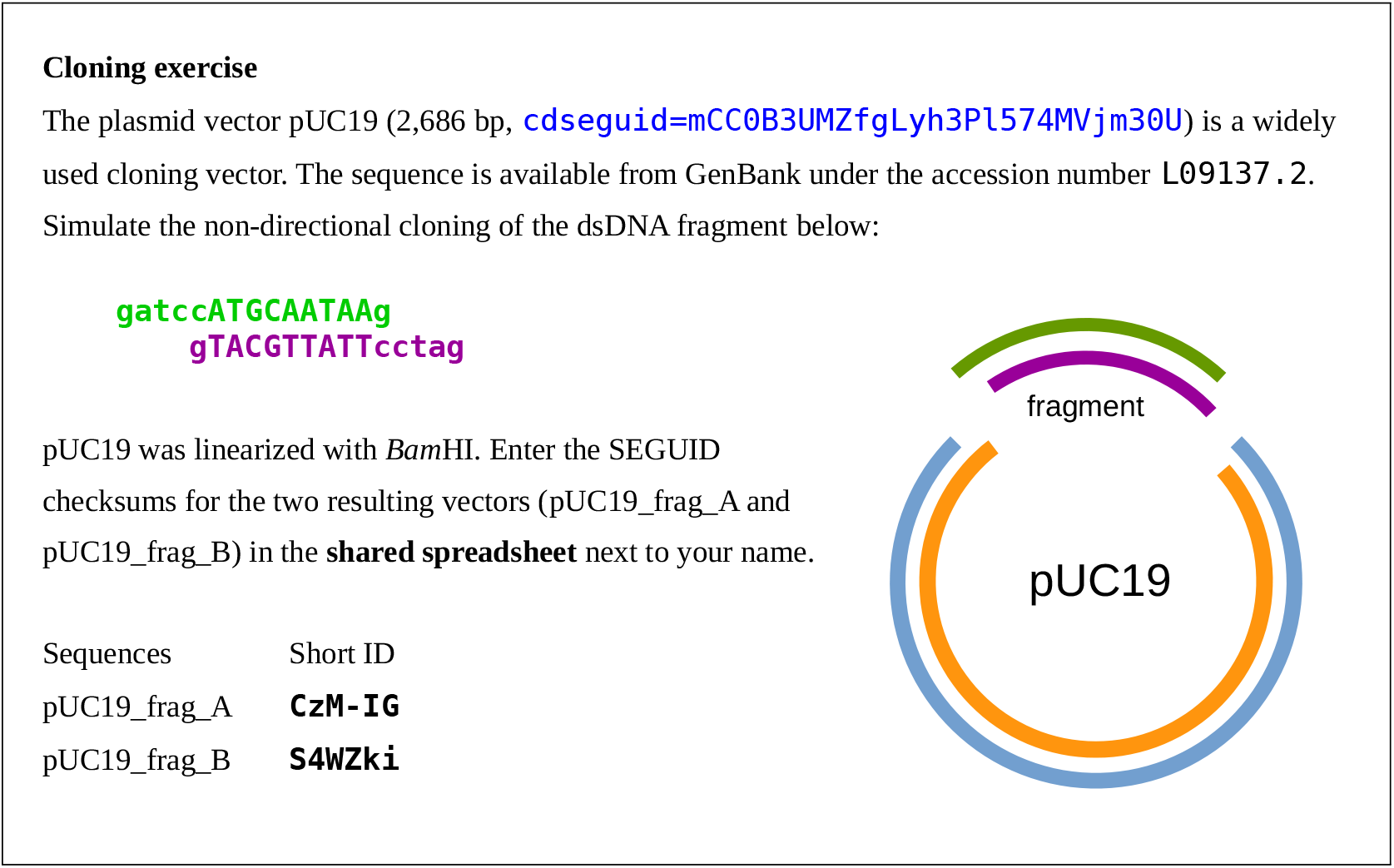
Cloning exercise. Students, in undergraduate courses at the Department of Biology of the University of Minho, are given a cloning simulation problem (top) that will result in two vectors. They are given the Short IDs of the correct vectors for them to self-correct before filing the results.

## 6 Discussions

The SEGUID v2 method produces checksums for various types of biological sequences, including, but not limited to, DNA, RNA, and proteins. The method is invariant to orientation of double-stranded sequences, and invariant to rotation for circular sequences. In contrast to other methods (e.g. Bouras et al. (2024)), SEGUID v2 is also independent of conventions or presence of specific sequences and does not require the input to comprise known gene sequences.

The SEGUID v2 checksum is capable of providing the scientific community with stable keys for the many thousands of plasmids created in the process of designing new recombinant organisms or synthetic biological constructs. Important plasmid sequences can be stored in a distributed fashion, without the need for expensive centralized infrastructure. SEGUID offers a stable, unique checksum for any DNA molecule, regardless of strandedness (single or double). The stability relies on the correctness of the underlying sequence. This means that sequence updates have to be accommodated in the long-term management of prior checksum versions for a sequence. The integrity of sequences distributed among members of the scientific community can easily be verified. Using SEGUID v2 makes it trivial to confirm that members of a research group are all using the same sequence for a particular DNA fragment containing a genetic part. The SEGUID v2 checksum would also enable a global reference service for circular DNA, such as the registry provided by EMBL-EBI at https://identifiers.org. Since the checksum is already implemented in the widely used DNA editor ApE, we expect this to encourage broader adoption among both researchers and web-facing applications. Future plans include continued maintenance and activities to drive wider use of the checksums.

## Acknowledgments

We would like to express our gratitude to the undergraduate students of Applied Biology and Biological Engineering at the University of Minho who have helped us to develop this work over the years.

BJ was funded by national funds via Fundação para a Ciência e Tecnologia Portugal (FCT) I.P. through project FatVal PTDC/EAM-AMB/032506/2017, and by the European Regional Development Fund (ERDF) via COMPETE2020 – Programa Operacional Competitividade e Internacionalização (POCI). The Center of Molecular and Environmental Biology Engineering (CBMA) was supported by the strategic program UIDB/04050/2020, funded by national funds via FCT I.P. HB was funded through the UCSF Computational Biology and Informatics (CBI) core that is supported by a National Cancer Institute (NCI) Cancer Center Support Grant 5P30CA082103. MWD was supported by NIH/NIGMS grant 1R01GM146005 (“Genome Engineering in the nematode C. elegans”). GB was supported through Argonne National Laboratory managed by UChicago Argonne, LLC for DOE under contract DE-AC02-06CH11357. This program is supported by the U.S. Department of Energy, Office of Science, through the Biomolecular Characterization and Sciences Program, Office of Biological and Environmental Research, under FWP 39156.

## Footnotes

1 https://www.uniprot.org/help/checksum

2 https://www.uniprot.org/help/uniparc

3 UniParc accession number *UPI0000117F35*_UP_

4 UniParc accession number *UPI0000117F39*

5 Strictly speaking, “smaller or equal” is sufficient to get a unique representation.

6 Technically, hash functions operate on raw bytes. To calculate a hash for a string, we ensure to use ASCII strings such that each character uniquely maps to an integer byte.

7 Technically, **Base64** encoding requires a multiple of four symbols. To meet this requirement, the 27-character string representing a **SHA-1** hash should be padded with a special symbol (=) at the very end.

8 The Short ID holds 6 symbols · log_2_ 64 bits/symbol = 36 bits (22.5%) of the original 160 bits of the **SHA-1** hash.

9 https://jorgensen.biology.utah.edu/wayned/ape

10 https://www.ncbi.nlm.nih.gov/genbank/samplerecord/

## Notes

https://www.seguid.org

### Competing Interest Statement

The authors have declared no competing interest.

### Summary of Updates

* Spell and grammar corrections * Minor rephrasing of sentences * Accession IDs corrections * Added minor discussions on SEGUID stability, sequence updates, ApE adoption

